# SARS-CoV-2 Nsp1 suppresses host but not viral translation through a bipartite mechanism

**DOI:** 10.1101/2020.09.18.302901

**Authors:** Ming Shi, Longfei Wang, Pietro Fontana, Setu Vora, Ying Zhang, Tian-Min Fu, Judy Lieberman, Hao Wu

## Abstract

The Severe Acute Respiratory Syndrome Coronavirus 2 (SARS-CoV-2) is a highly contagious virus that underlies the current COVID-19 pandemic. SARS-CoV-2 is thought to disable various features of host immunity and cellular defense. The SARS-CoV-2 nonstructural protein 1 (Nsp1) is known to inhibit host protein translation and could be a target for antiviral therapy against COVID-19. However, how SARS-CoV-2 circumvents this translational blockage for the production of its own proteins is an open question. Here, we report a bipartite mechanism of SARS-CoV-2 Nsp1 which operates by: (1) hijacking the host ribosome via direct interaction of its C-terminal domain (CT) with the 40S ribosomal subunit and (2) specifically lifting this inhibition for SARS-CoV-2 via a direct interaction of its N-terminal domain (NT) with the 5’ untranslated region (5’ UTR) of SARS-CoV-2 mRNA. We show that while Nsp1-CT is sufficient for binding to 40S and inhibition of host protein translation, the 5’ UTR of SARS-CoV-2 mRNA removes this inhibition by binding to Nsp1-NT, suggesting that the Nsp1-NT-UTR interaction is incompatible with the Nsp1-CT-40S interaction. Indeed, lengthening the linker between Nsp1-NT and Nsp1-CT of Nsp1 progressively reduced the ability of SARS-CoV-2 5’ UTR to escape the translational inhibition, supporting that the incompatibility is likely steric in nature. The short SL1 region of the 5’ UTR is required for viral mRNA translation in the presence of Nsp1. Thus, our data provide a comprehensive view on how Nsp1 switches infected cells from host mRNA translation to SARS-CoV-2 mRNA translation, and that Nsp1 and 5’ UTR may be targeted for anti-COVID-19 therapeutics.

## INTRODUCTION

Severe Acute Respiratory Syndrome Coronavirus 2 (SARS-CoV-2), the causative agent of the newly emerged infectious disease COVID-19, is a highly contagious and deadly virus with fast person-to-person transmission and potent pathogenicity (Chan et al., 2020b). It is an enveloped, single-stranded betacoronavirus that contains a positive-sense RNA genome of about 29.9 kb (Lu et al., 2020; Wang et al., 2020; Wu et al., 2020; Zhu et al., 2020). The SARS-CoV-2 genome codes for two large overlapping open reading frames in gene 1 (ORF1a and ORF1b) and a variety of structural and nonstructural accessory proteins (Zhou et al., 2020). Upon infection, SARS-CoV-2 hijacks host cell translation machinery to synthesize ORF1a and ORF1b polyproteins that are subsequently proteolytically cleaved into 16 mature non-structural proteins, namely Nsp1 to Nsp16 (Chan et al., 2020a; Hartenian et al., 2020; Zhou et al., 2020).

Nsp1 is a critical virulence factor of coronaviruses and plays key roles in suppressing host gene expression, which facilitates viral replication and immune evasion (Jimenez-Guardeno et al., 2015; Kamitani et al., 2009; Tanaka et al., 2012). It has been shown that Nsp1 effectively suppresses the translation of host mRNAs by directly binding to the 40S small ribosomal subunit (Narayanan et al., 2008; Tohya et al., 2009). The recent cryo-electron microscopy (cryo-EM) structures indeed revealed the binding of the C-terminal domain (CT) of Nsp1 to the mRNA entry channel of the 40S (Schubert et al., 2020; Thoms et al., 2020), blocking translation. However, how SARS-CoV-2 overcomes the Nsp1-mediated translation suppression for its own replication remains an open question.

Previous studies on SARS-CoV suggest that the the 5’ untranslated region (5’ UTR) of coronavirus mRNA protects the virus against Nsp1-mediated mRNA translation inhibition (Huang et al., 2011; Kamitani et al., 2009; Thoms et al., 2020). Here, we investigated the mechanism of how Nsp1 suppresses translation and how SARS-CoV-2 escapes this suppression to effectively switch the translational machinery from synthesizing host proteins to making viral proteins during its infection.

## RESULTS

### SARS-CoV-2 Nsp1 Suppresses Translation

To investigate the function of SARS-CoV-2 Nsp1 in inhibiting mRNA translation, we co-transfected an mScarlet reporter construct with MBP-tagged Nsp1 or MBP alone in HeLa cells and imaged mScarlet fluorescence and anti-MBP immunofluorescence (Figure 1A). The mScarlet reporter used an expression vector that contains the cytomegalovirus (CMV) promoter and 5’ UTR (referred as control 5’ UTR) and is commonly employed for mammalian cell expression. Compared to cells transfected with the MBP control, MBP-Nsp1-transfected cells showed markedly reduced mScarlet expression. Quantification of mScarlet reporter intensities showed that MBP-Nsp1 decreased mScarlet expression by over 4-fold (*p* < 0.001) (Figure 1B). These data demonstrate that SARS-CoV-2 Nsp1 potently inhibits protein translation in cells, recapitulating previous reports (Narayanan et al., 2008; Schubert et al., 2020; Thoms et al., 2020; Tohya et al., 2009).

**Figure 1.**
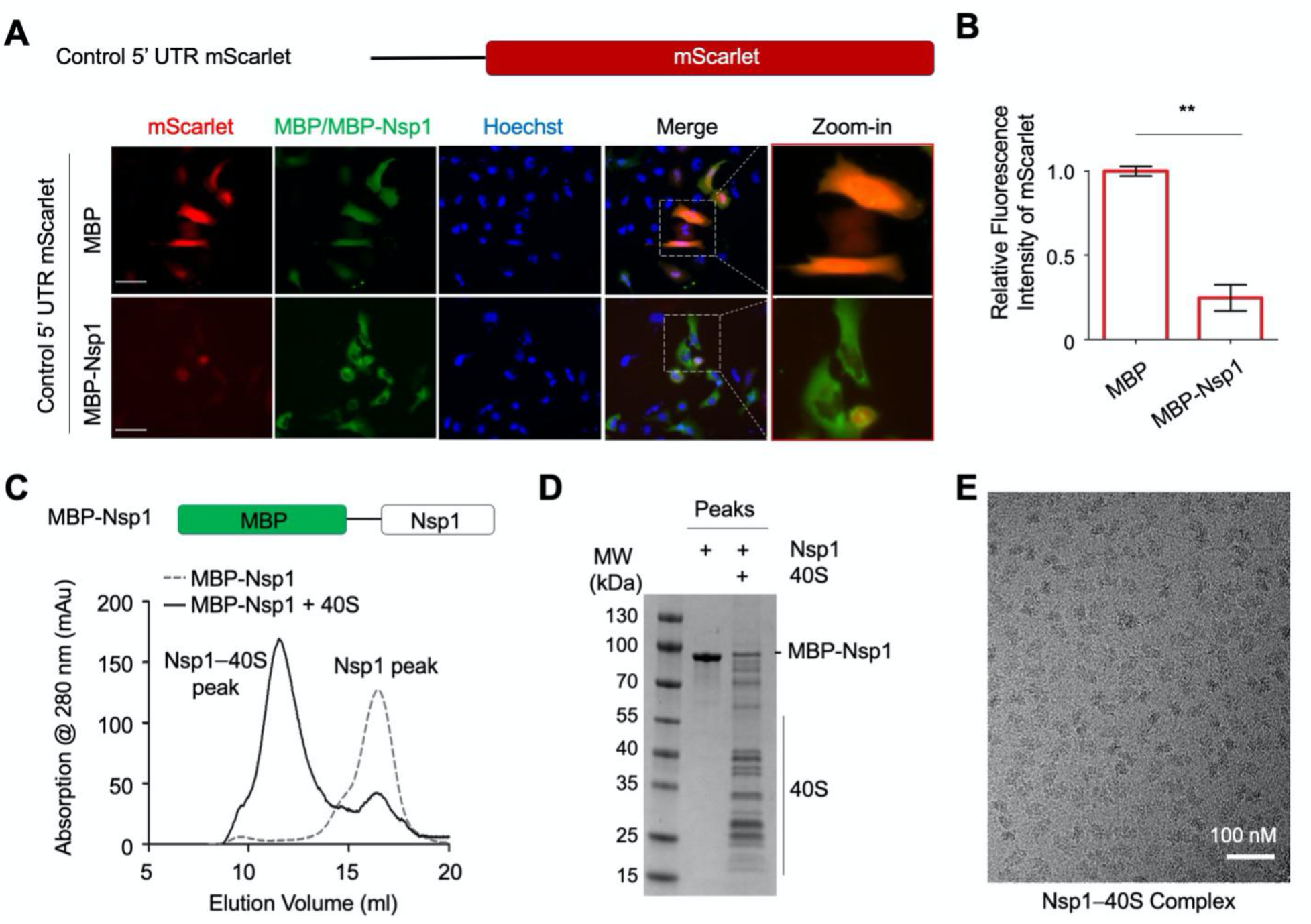
Suppression of Protein Synthesis by SARS-CoV-2 Nsp1. (A) Schematic representation of CMV 5’ UTR mScarlet reporter (upper panel) and immunofluorescence images of HeLa cells transfected with mScarlet (red) and MBP or MBP-Nsp1 (green), and DNA (blue) (lower panel). (B) Quantification of relative fluorescence intensity of mScarlet in MBP- or MBP-Nsp1-expressing HeLa cells. (C) Gel filtration profiles of MBP-Nsp1 and the MBP-Nsp1–40S complex. Schematic representation of MBP-Nsp1 is shown. Peak positions are labeled. (D) SDS PAGE of MBP-Nsp1 and the MBP-Nsp1–40S complex. (E) A cryo-EM micrograph of the MBP-Nsp1–40S complex.

### The C-terminal Helical Hairpin of Nsp1 Binds to the mRNA Channel of 40S Ribosome

To elucidate how Nsp1 inhibits protein translation, we set out to determine the structure of an Nsp1–40S complex using cryo-EM. We first expressed and purified His-MBP-tagged Nsp1 by Ni-NTA affinity and gel filtration chromatography (Figure 1C). We then purified human ribosomes from Expi293 cells using an established protocol (Khatter et al., 2014), dissociated the large and small subunits (60S and 40S) by puromycin treatment, and further purified 60S and 40S away from mRNAs and other contaminants by sucrose cushion. Mixing the ribosome preparation with MBP-Nsp1 followed by amylose affinity and gel filtration chromatography yielded the Nsp1–40S complex (Figure 1C-D). The sample displayed homogenous particles on cryo-EM micrographs (Figure 1E).

We determined the cryo-EM structure of the Nsp1–40S complex at a resolution of at least 2.9 Å for both the body and the head regions of the 40S (Figure 2A, S1). The 40S structure was rigidly fit using the PDB entry 6g5h (Ameismeier et al., 2018). The Nsp1-CT could be traced in the cryoEM map, which revealed a tight helical hairpin (Figure 2A-C). However, the MBP tag and the N-terminal domain (Nsp1-NT) are invisible in the cryo-EM map, suggesting that they are flexibly linked to the CT. Consistently, there is a predicted ~20 residue linker between Nsp1-NT and Nsp1-CT. Nsp1 interacts with the 40S at the mRNA channel located at the interface between the head and body of the 40S (Figure 2B). As such, Nsp1 binding blocks mRNA entry to the ribosome to inhibit protein translation, in agreement with recent structural studies (Schubert et al., 2020; Thoms et al., 2020).

**Figure 2.**
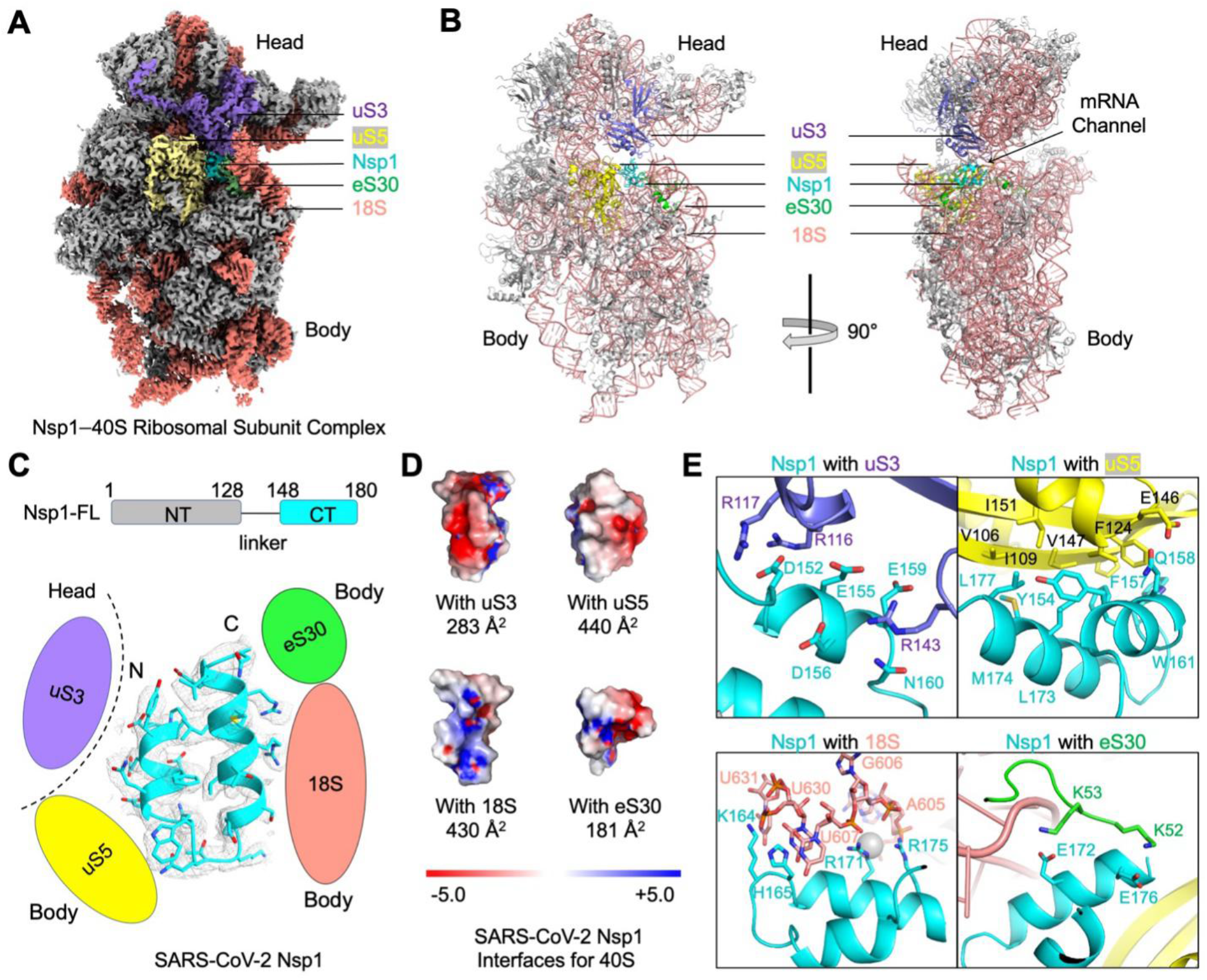
Blockage of the mRNA Channel in the 40S by the Nsp1 C-terminal Helical Hairpin. (A) Cryo-EM density of the Nsp1–40S complex. Nsp1 is in cyan. Subunits of 40S that interact with Nsp1 are color-coded. (B) Ribbon diagram of the Nsp1–40S complex. Nsp1 (cyan) binds at the mRNA channel in the cleft between the head and body of the 40S ribosomal subunit. Subunits of 40S that interact with Nsp1 are color-coded. (C) Ribbon diagram and cryo-EM density of SARS-CoV-2 Nsp1 with schematics highlighting its interfaces to 40S. (D) Electrostatic surface representations of Nsp1 binding surfaces to the 40S subunits and rRNA. Buried surface areas are marked. (E) Detailed interactions between Nsp1 and the 40S ribosomal subunit. See also Supplemental Figure 1

In more detail, Nsp1 forms interactions with three 40S protein subunits, uS3, uS5 and eS30, as well as the 18S rRNA (Figure 2B-C). It is worth noting that three out of the four binding interfaces are between Nsp1 and the body of the 40S (uS5, eS30, 18S); only one interface, at the most proximal region of the Nsp1-CT, is with the head of the 40S (uS3) (Figure 2D). The Nsp1-uS3 interface has a buried surface area of 283 Å^2^ with mostly charge-charge interactions, and is mediated by Nsp1 residues, including D152, E155, E159 and D156 (Figure 2E). In contrast, the Nsp1-uS5 interface has a larger buried surface area of 440 Å^2^, with mainly hydrophobic interactions (Figure 2E). At the interface with the18S rRNA, Nsp1 residues H165 and K164 stack with the bases of U607, U630, and U631, while R171 and R175 form charge-charge interactions with the phosphate backbone of A604, G606 and U607 (Figure 2E). For the Nsp1–eS30 interaction, negatively charged residues E172 and E176 of Nsp1 interact with positively charged residues K52 and K53 at one extended loop of eS30 (Figure 2E). Overall, Nsp1 is highly buried, with close to 60% of its total surface area of 2,400 Å^2^ interacting with the 40S.

We hypothesized that the single interface of Nsp1 with the 40S head is likely cannot restrict the rotation and the intrinsic flexibility of the 40S head relative to the 40S body, which is known to be necessary during protein translation (Hussain et al., 2014; Lomakin and Steitz, 2013). Consistent with this hypothesis, we observed certain degrees of movement between the head and body of the 40S during 3D classification and reconstruction; as a result, high-resolution cryo-EM maps were obtained using focused refinement (Figure S1). We postulate that the flexibility of the head, as well as its limited interaction with Nsp1-CT, may be important for the ability of SARS-CoV-2 5’ UTR to dislodge the Nsp1–40S interaction to evade the translation block (see DISCUSSION).

### The 5’ UTR of SARS-CoV-2 mRNA Bypasses the Translation Inhibition by Nsp1

To analyze the effect of SARS-CoV-2 Nsp1 on the translation of SARS-CoV-2 mRNA, we replaced the control 5’ UTR in the mScarlet reporter construct with the SARS-CoV-2 5’ UTR (Figure 3A), and co-expressed it with MBP-tagged Nsp1 or MBP alone in HeLa cells (Figure 3B-C). In contrast to the control 5’ UTR, expression of the mScarlet reporter downstream from SARS-CoV-2 5’ UTR in MBP-Nsp1 transfected cells was not significantly different from MBP transfected control cells. As expected, MBP-Nsp1-NT, which does not interact with the 40S, also did not affect mScarlet reporter expression. By contrast, a severe reduction in mScarlet expression was observed by MBP-Nsp1-CT (*p* < 0.001). Co-expression of MBP-Nsp1-NT did not remove the inhibitory effect of MBP-Nsp1-CT on mScarlet expression, suggesting that only covalently linked NT and CT could rescue mScarlet expression downstream the SARS-CoV-2 5’ UTR. These results demonstrate that full-length Nsp1 selectively suppresses host but not SARS-CoV-2 protein translation, while the Nsp1-CT alone is a general inhibitor of both host and viral protein synthesis.

**Figure 3.**
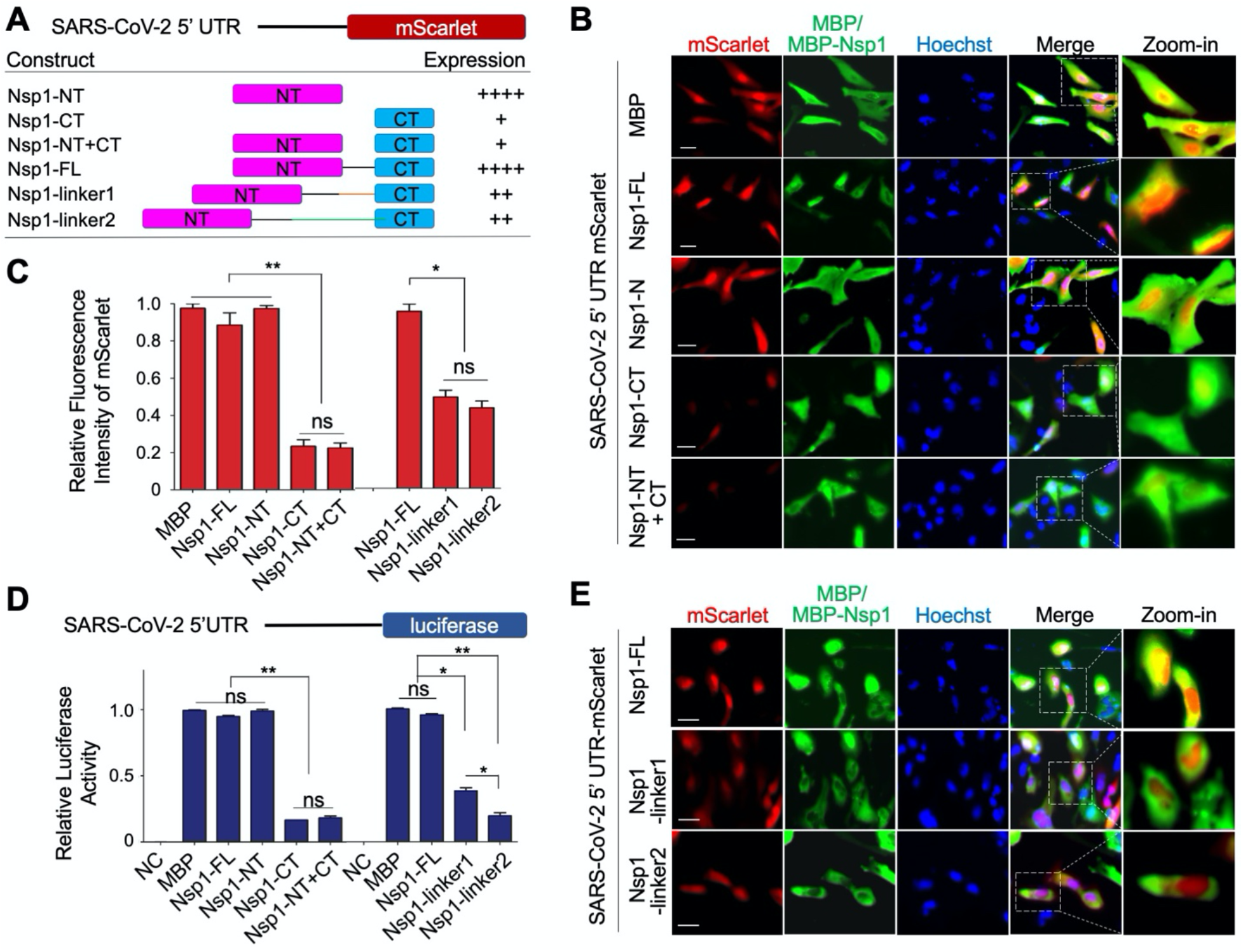
Evasion of Nsp1-Mediated Translation Inhibition by SARS-CoV-2 5’ UTR. (A) Summary of Nsp1 constructs and SARS-CoV-2 5’ UTR mScarlet reporter levels. The various Nsp1 constructs are: Nsp1-FL, Nsp1-NT, Nsp1-CT, both Nsp1-NT and Nsp1-CT (Nsp1-NT+CT), Nsp1-linker1, and Nsp1-linker2. (B) Immunofluorescence images of HeLa cells transfected with SARS-CoV-2 5’ UTR mScarlet reporter (red) and MBP, MBP-Nsp1-FL, MBP-Nsp1-NT, MBP-Nsp1-CT, or MBP-Nsp1-N+C (green). (C) Quantification of relative fluorescence intensity of mScarlet in indicated groups from (B) and (E). (D) Quantification of the relative luciferase activity in HEK293T cells transfected with SARS-CoV-2 5’ UTR luciferase reporter and various Nsp1 constructs. Values are means ± S.E.M. obtained from three independent experiments. (E) Immunofluorescence images of HeLa cells transfected with SARS-CoV-2 5’ UTR mScarlet reporter (red) and Nsp1-FL, Nsp1-linker1, or Nsp1-linker2 (green).

Next, we introduced a SARS-CoV-2 5’ UTR luciferase reporter in HEK293T cells to confirm the observations derived from the mScarlet report assay (Figure 3D). Consistently, MBP-Nsp1 and Nsp1-NT did not inhibit the translation of SARS-CoV-2 5’ UTR-luciferase, whereas both Nsp1-CT and Nsp1-NT + CT displayed indistinguishably decreased translation.

Because covalently linking Nsp1-NT and CT is required for the evasion of Nsp1-mediated translation inhibition by SARS-CoV-2 5’ UTR, we wondered if the length of the linker between NT and CT in Nsp1 has any functional effect. To address this question, we inserted additional 20 residues (linker1) or 40 residues (linker2) at the Nsp1 linker (Figure 3A). Remarkably, the linker extension dramatically reduced the ability of SARS-CoV-2 5’ UTR to evade Nsp1-mediated translation inhibition as measured by mScarlet expression (*p* < 0.05) (Figure 3C, E). These data were further validated in the SARS-CoV-2 5’ UTR luciferase assay. Interestingly, compared with the 20-residue linker, the 40-residue linker was more effective in suppressing SARS-CoV-2 5’ UTR-mediated evasion of translation inhibition (Figure 3D).

### SARS-CoV-2 5’ UTR Directly Interacts with Nsp1-NT

Since Nsp1-CT is a general inhibitor of mRNA translation, clarifying the explicit function of Nsp1-NT is important for understanding how SARS-CoV-2 5’ UTR evades inhibition mediated by Nsp1. We investigated whether SARS-CoV-2 5’ UTR directly interacts with Nsp1-NT. For this purpose, we generated a construct with strep-tagged Nsp1-NT downstream of SARS-CoV-2 5’ UTR and expressed it in Expi293F cells (Figure 4A). Upon affinity purification using the strep tag followed by gel filtration chromatography, we immunoblotted the peak fractions using anti-strep to detect Nsp1-NT, and performed RT-PCR on the same fractions to detect SARS-CoV-2 5’ UTR. Notably, Nsp1-NT and SARS-CoV-2 5’ UTR co-eluted in the same peak (Figure 4B), suggesting a direct interaction between them.

**Figure 4.**
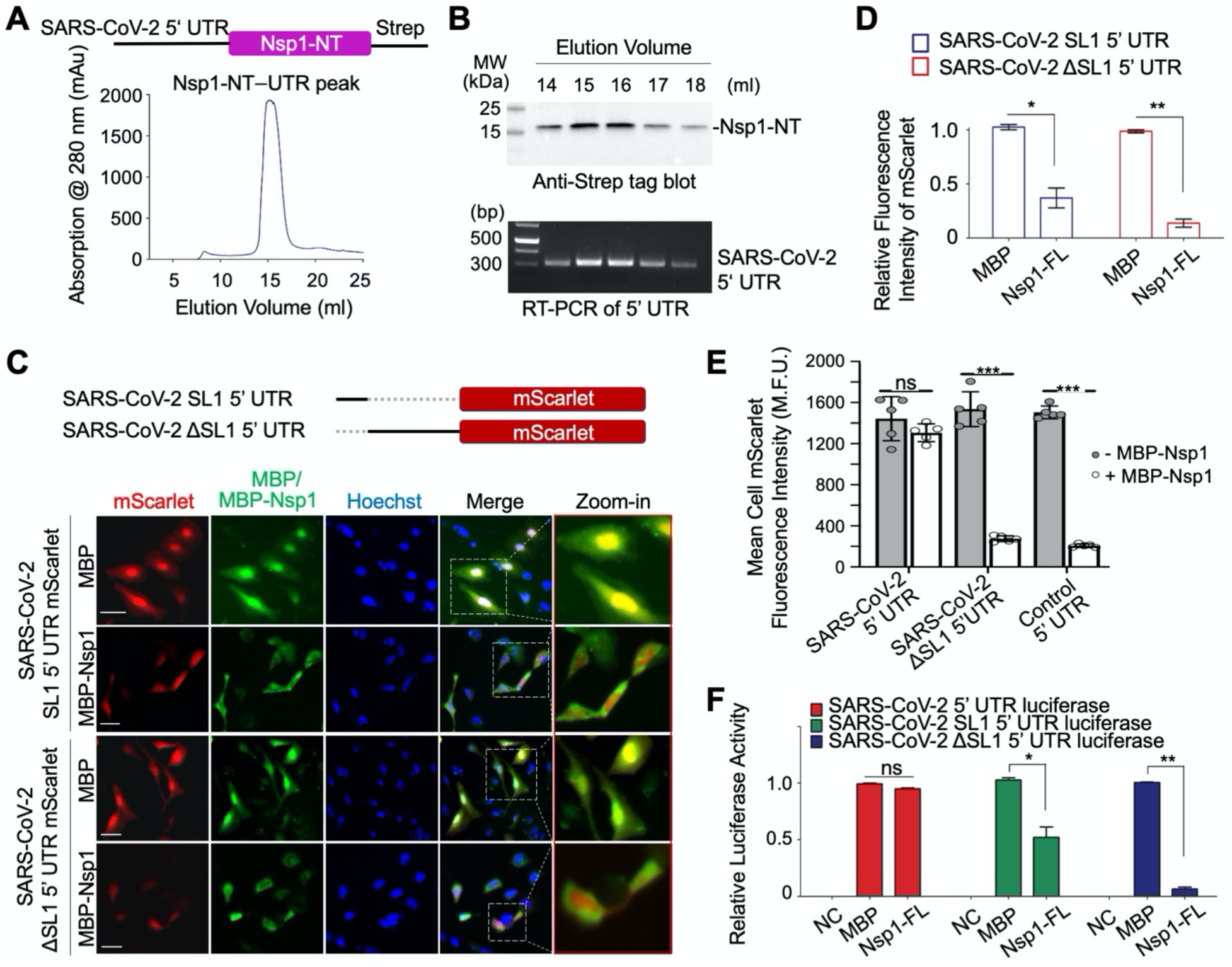
Direct interaction between SARS-CoV-2 5’ UTR and Nsp1-NT, and Requirement of the SL1 of the 5’ UTR in Viral Evasion. (A) Gel filtration profile of the complex between Nsp1-NT (Strep-tagged) and SARS-CoV-2 5’ UTR purified by the Strep-Tactin^®^ affinity resin from the HeLa cell lysate transfected with the SARS-CoV-2 5’ UTR Nsp1-NT-Strep construct. (B) Anti-Strep tag Western blot (upper panel) and RT-PCR of SARS-CoV-2 5’ UTR (lower panel) of the gel filtration peak fractions in (A). Nsp1-NT co-migrated with SARS-CoV-2 5’ UTR. (C) Construct design of SARS-CoV-2 SL1 5’ UTR and SARS-CoV-2 ΔSL1 5’ UTR-mScarlet reporters (upper panel) and immunofluorescence images of HeLa cells transfected with a mScarlet reporter (red) and MBP or MBP-Nsp1-FL (green) (lower panel) (D) Quantification of data in (C) showing the relative fluorescence intensity of mScarlet in HeLa cells transfected with MBP or MBP-Nsp1-FL. (E) High throughput quantification of mean cellular mScarlet fluorescence intensity when placed downstream of either SARS-CoV-2 5’ UTR, SARS-CoV-2 ΔSL1 5’ UTR, or control 5’UTR, with or without co-expression of MBP-Nsp1. (F) Relative luciferase activity in HEK293T cells after co-transfection of a luciferase reporter and MBP or MBP-Nsp1. The luciferase reporter was placed downstream of SARS-CoV-2 5’ UTR, SARS-CoV-2 SL1 5’ UTR or SARS-CoV-2 ΔSL1 5’ UTR. NC: negative control without adding the luciferase substrate in cells co-transfected with luciferase reporter and MBP. Values are means ± S.E.M. obtained from three independent experiments.

### The SL1 Stem-Loop of the 5’ UTR Is Required for the Translation of SARS-CoV-2 mRNA

The 5’ UTR of coronaviruses comprises at least four stem-loop structures, among which stemloop 1 (SL1) has been shown to play critical roles in driving viral replication (Li et al., 2008). Thus we generated SARS-CoV-2 SL1 and ΔSL1 5’ UTR mScarlet reporters to elucidate the function of SL1. Compared with mScarlet translation in control cells co-transfected with MBP, the translation of SARS-CoV-2 ΔSL1 5’ UTR mScarlet reporter was markedly reduced in MBP-Nsp1 transfected cells (*p* < 0.001), suggesting that SL1 is required for evasion of Nsp1-mediated translation suppression (Figure 4C-D). In comparison with SARS-CoV-2 ΔSL1 5’ UTR, the 5’ UTR containing only the SL1 region retained more mScarlet expression, suggesting that SL1 is partially active against the translation block by Nsp1; however, SL1-mediated mScarlet expression is significantly less in Nsp1-transfected cells than in MBP-transfected cells, suggesting that SL1 alone is not sufficient to evade the translation suppression (Figure 4C-D).

The role of SL1 was further confirmed by high throughout analysis of mScarlet-expressing cells using an ImageXpress high content microscope. In contrast to the intact SARS-CoV-2 5’ UTR, which showed no change in reporter activity in the presence or absence of Nsp1, cells expressing the SARS-CoV-2 ΔSL1 5’ UTR reporter showed dramatically reduced mScarlet mean fluorescence intensity upon Nsp1 co-expression (Figure 4E). We also generated SARS-CoV-2 SL1 or ΔSL1 5’ UTR luciferase reporter and found that SL1 conferred partial antagonism to translation suppression by Nsp1, while ΔSL1 lost the ability to escape the translation suppression (Figure 4F). Together, these data support the role of SL1 in SARS-CoV-2 5’ UTR as an important element in the evasion of translation inhibition. Curiously, a previous study failed to show SARS-CoV-2 5’ UTR-mediated evasion of Nsp1-mediated translation inhibition (Schubert et al., 2020). However, judging from the primer used, the 5’ UTR in the experiment did not contain SL1, consistent with our data on SARS-CoV-2 ΔSL1 5’ UTR.

## DISCUSSION

SARS-CoV-2 Nsp1 is a major virulence factor which suppresses host gene expression and immune defense (Jimenez-Guardeno et al., 2015; Kamitani et al., 2009; Tanaka et al., 2012). Consistent with recently published cryo-EM structures (Schubert et al., 2020; Thoms et al., 2020), our data showed that the helical hairpin at the CT of SARS-CoV-2 Nsp1 interacts with the 40S subunit of the ribosome to block mRNA entry (Figure 5), confirming that Nsp1-CT is a general inhibitor of the cellular translation machinery. More importantly, we revealed that Nsp1-NT directly interacts with SARS-CoV-2 5’ UTR, and that this interaction relieves translational inhibition imposed by Nsp1-CT to allow successful translation of the SARS-CoV-2 mRNA (Figure 5). Thus, coronaviruses like SARS-CoV-2 have evolved a clever strategy to selectively hijack the host translation machinery to support their own replication while counteracting the host immune response.

**Figure 5.**
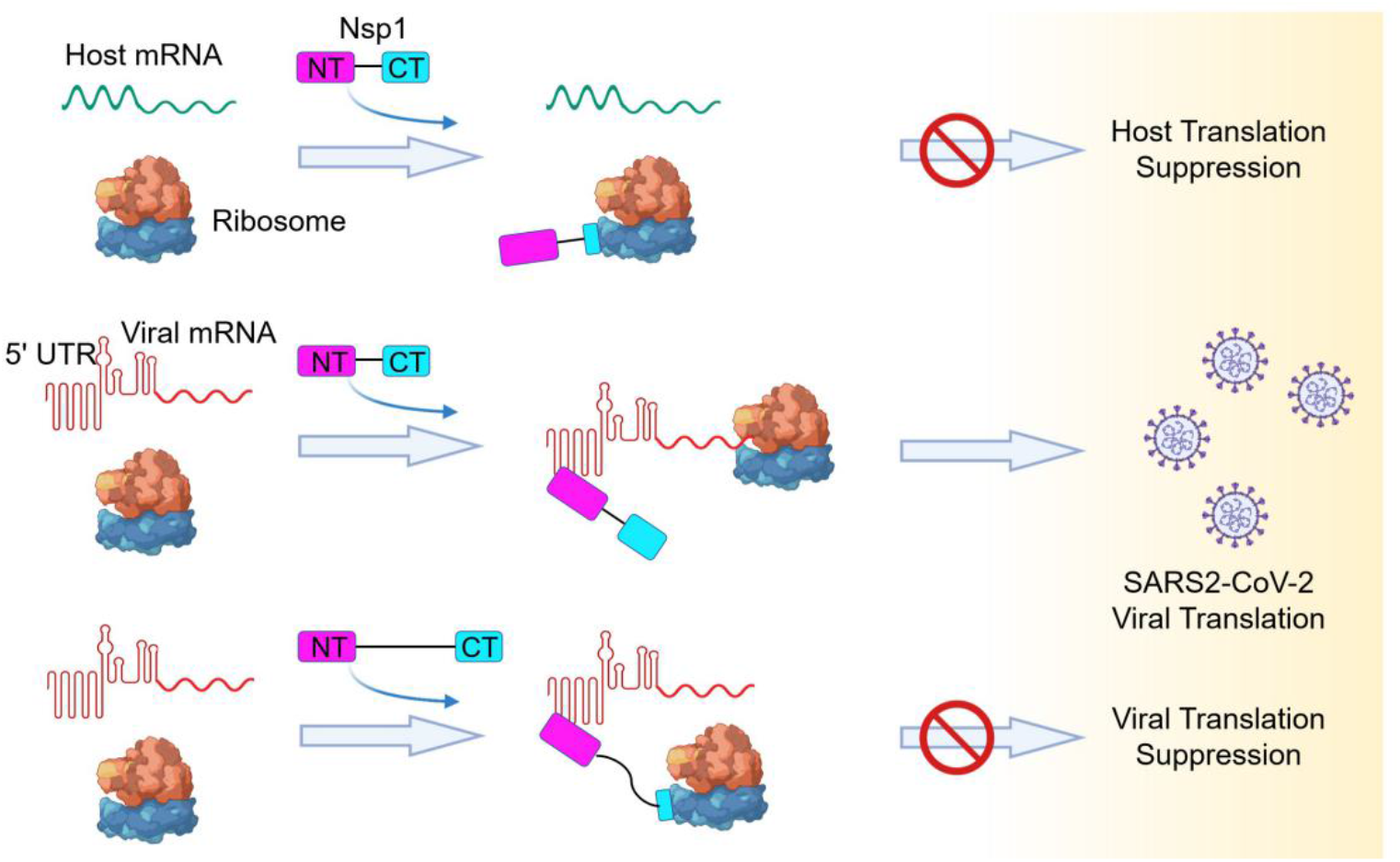
Model of a Bipartite Mechanism for Nsp1-Mediated Translation Inhibition and Evasion by SARS-CoV-2 5’ UTR. A schematic model depicting the bipartite roles of SARS-CoV-2 Nsp1 during infection. First, Nsp1 blocks host mRNA from binding to the 40S ribosomal subunit due to physical occlusion by the bound Nsp1-CT. Second, Nsp1 supports viral mRNA translation by interacting with SARS-CoV-2 5’ UTR using Nsp1-NT, which results in dissociation of the Nsp1-CT–40S complex to overcome inhibition. This mechanism of evasion of Nsp1-mediated translation inhibition is illustrated by the failure of linker-lengthened Nsp1 to support viral mRNA translation. With the longer linker, the Nsp1-NT–5’ UTR complex can co-exist with the Nsp1-CT–40S complex.

We hypothesize that the Nsp1-CT-40S interaction is incompatible with the interaction between SARS-CoV-2 5’ UTR and Nsp1-NT. When SARS-CoV-2 5’ UTR is bound to Nsp1-NT, the covalently linked Nsp1-CT cannot bind 40S likely due to a steric factor in which Nsp1-CT is unable to reach 40S for a productive interaction. This in turn allows loading of the SARS-CoV-2 mRNA for translation initiation. Consistent with this hypothesis, increasing the linker length between Nsp1-NT and Nsp1-CT greatly reduced SARS-CoV-2 5’ UTR-mediated translation in the presence of Nsp1 (Figure 5). Even when an Nsp1 molecule is already bound to a ribosome, we postulate that the SARS-CoV-2 5’ UTR can interact with Nsp1-NT to strip off the Nsp1-CT-40S interaction to proceed with viral mRNA translation. One possible scenario is that the binding of SARS-CoV-2 5’ UTR to Nsp1-NT exerts a pulling force via the short linker to the N-terminus of Nsp1-CT, which is weakly bound at the flexible head of the 40S, to dislodge Nsp1-CT from the 40S. This mechanism may be analogous to how newly synthesized IκBα can remove the NF-κB transcription factor from the bound DNA (Oeckinghaus and Ghosh, 2009).

Thus, there is a tug of war between Nsp1-NT and Nsp1-CT, pulled by their respective binding partners, SARS-CoV-2 5’ UTR and the 40S subunit of the ribosome. Our data reveal the importance of the SL1 stem-loop at the beginning of SARS-CoV-2 5’ UTR in overcoming the inhibition of translation by Nsp1 and highlight a new potential therapeutic target for limiting SARS-CoV-2 replication. Blocking the 5’ UTR-Nsp1 interaction, be it by a small molecule, an anti-sense RNA or other mechanisms, may resolve this tug of war to the detriment of the SARS-CoV-2 virus to help treat the devastating COVID-19.

## EXPERIMENTAL PROCEDURES

### Plasmids, Cell Culture and Transfection

SARS-CoV-2 full-length Nsp1 (1-180 aa), Nsp1-NT (1-127 aa) and Nsp1-CT (128-180 aa) were amplified from pDONR207 SARS-CoV-2 NSP1 (Addgene) by PCR and then cloned into pDB-His-MBP or BacMam pCMV-Dest plasmid. Full-length 265 nt 5’ UTR of SARS-CoV-2 was subcloned to replace the 5’ UTR of human CMV in the pLV-mScarlet vector using a Hifi one-step kit (Gibson Assembly, NEB). Full-length, SL1 alone or ΔSL1 5’ UTR of SARS-CoV-2 were cloned into pLV-mScarlet vector or pGL3 basic vector. All constructs were verified by sequencing.

HEK293T, Expi293F and HeLa cells were purchased from the American Type Culture Collection (ATCC), and cultured in Dulbecco’s Modified Eagle medium (DMEM) or Expi293 Expression Medium (Gibco) supplemented with 10% fetal bovine serum and 1% penicillin/streptomycin (Gibco) at 37 °C with 5% CO_2_. Cells were transiently transfected with indicated plasmids using FuGENE Transfection Reagent (Promega) or Lipofectamine 2000 (Thermo Fischer Scientific) according to the manufacturer’s instructions.

### Preparation of Nsp1, Human Ribosome and the Nsp1-40S Complex

SARS-CoV-2 full-length Nsp1 (1-180 aa), Nsp1-NT (1-127 aa) and Nsp1-CT (128-180 aa) constructs carrying an N-terminal His-MBP tag were transformed into *E. coli* BL21-CodonPlus (DE3)-RIPL, and single colonies were picked and grown in LB medium supplemented with 50 μg/ml kanamycin at 37 °C. Protein expression was induced for 16 h at 18 °C with 0.5 mM IPTG after OD_600_ of the culture reached 0.8. Cells were harvested by centrifugation, resuspended in lysis buffer (20 mM Tris pH 7.4, 150 mM NaCl, 1 mM TCEP and protease inhibitors) and lysed using ultrasonic homogenizer (Constant Systems Ltd). The proteins were first purified by affinity chromatography using TALON metal affinity resin (TaKaRa), and further purified via gel filtration chromatography on a Superdex 200 column (GE Healthcare) in gel filtration buffer (20 mM Tris pH 7.4, 150 mM NaCl and 1 mM TCEP). The purified proteins were flash-frozen in liquid nitrogen, and stored until further use at −80 °C.

Human 80S ribosome was purified as described (Khatter et al., 2014), followed by incubation with 1 mM puromycin for 30 min at 4 °C in buffer R containing 20 mM Tris pH 7.5, 2 mM MgCl_2_, 150 mM KCl, and 1 mM TCEP. The mixture containing 40S and 60S subunits was loaded onto a 30% sucrose cushion and centrifuged to remove mRNAs and other contaminants. The pellets were resuspended in buffer R, flash-frozen in liquid nitrogen, and stored at −80 °C. Nsp1-ribosome complex formation was carried out by incubating recombinant His-MBP-Nsp1 protein with the ribosome preparation containing 40S and 60S subunits for 30 min. Nsp1–40S complex was then affinity-purified using amylose resin (NEB), followed by size-exclusion chromatography. The peak corresponding to the Nsp1–40S complex was collected, concentrated to 2 OD_280_, and subjected to cryo-EM analysis.

### Cryo-EM Data Collection and Processing

A 3 μl drop of the Nsp1–40S complex was applied to glow-discharged copper grids with lacey carbon support and a 3 nm continuous carbon film (Ted Pella, Inc). The grids were blotted for 4 s in 100% humidity at 4 °C, and plunged-frozen using an FEI Vitrobot Mark IV. Cryo-EM data collection was performed on a 300 keV Titan Krios microscope (FEI) with a K3 direct electron detector (Gatan) at the National Cancer Institute’s National Cryo-EM Facility. 16,119 movies were collected in counting mode, with 40 total frames per movie, 3.2 s exposure time, 50 electrons per Å^2^ accumulated dose, and 1.08 Å physical pixel size. Movies were motion-corrected and dose-weighted using MotionCor2 (Zheng et al., 2017). Patch contrast transfer function (CTF) estimation was performed using cryoSPARC (Punjani et al., 2017). Particle picking was initially carried out using blob picker in cryoSPARC, followed by neural network based particle picking using Topaz (Bepler et al., 2019). Particle extraction was carried out with a box size of 360 pixels, followed by 2D classification in cryoSPARC. Classes were manually selected. Particles from selected classes were then 3D classified into three classes using the three reference maps generated from ab-initio 3D reconstruction, resulting in one class with good orientation distribution. Particles from the selected 3D class were “polished” through a Bayesian polishing process in Relion (Zivanov et al., 2019). “Polished” particles were imported into cryoSPARC to perform non-uniform 3D refinement (Punjani et al., 2017), which gave a map at 2.86 Å resolution. Focused refinements of head and body regions of the 40S were performed using the local refinement in cryoSPARC, which resulted in a 2.86 Å map of the body region and a 2.89 Å map of the head region. The focused-refined maps were used to generate a composite map of the Nsp1–40S complex in UCSF Chimera (Pettersen et al., 2004).

The 40S subunit was modeled by rigid body fitting of a human 40S model (PDB ID:6g5h) (Ameismeier et al., 2018) into the cryo-EM maps using Chimera (Pettersen et al., 2004). Nsp1 was built de-novo based on the cryo-EM map. Inspection, model building, and manual adjustment were carried out in Coot (Emsley and Cowtan, 2004). Real-space refinement was performed using PHENIX (Adams et al., 2010). All representations of densities and structural models were generated using Chimera, ChimeraX (Goddard et al., 2018), and Pymol (DeLano, 2002).

### Semi-Quantitative Reverse Transcription-PCR and Western Blot Analysis

Reverse transcription-PCR (RT-PCR) was performed using the PrimeScript RT Reagent Kit (TaKaRa). Quantification of mRNA level was performed in a 20 μl mixture consisting of 10 μl Q5 High-Fidelity 2X Master Mix (NEB), 0.2 μl RT-PCR product, 1 μl primer set mix at a concentration of 5 pmol/ml for each primer and 8.8 μl sterile water. The forward primer of SARS-CoV-2 5’ UTR is 5’ ATTAAAGGTTTATACCTTCCCAG 3’, and the reverse primer is 5’ CTTACCTTTCGGTCACAC 3’. Cells were collected and lysed in RIPA lysis buffer (Thermo Scientific) with complete protease inhibitor cocktail (Roche Applied Science). The protein levels were determined using Western blot analysis. Equal portions of the cell lysate were run on a 15% SDS-PAGE gel and blotted onto a PVDF membrane, which was subsequently probed with HRP Anti-Strep tag antibody (Abcam) and developed with an ECL substrate (Amersham Biosciences).

### Luciferase Assay

Luciferase assays were performed as previously described (Yang et al., 2020). Briefly, HEK293T cells were transfected with control 5’ UTR, full-length, SL1 alone, or SL1-deleted SARS-CoV-2 5’ UTR luciferase reporter plasmid using FuGENE Transfection Reagent (Promega). Luciferase assays were performed using the Luciferase Assay System (Promega); β-Galactosidase activity was used as an internal control. Luciferase activity was measured using a CytoFluorplate 4000 Luminescence Microplate Reader (ABI).

### Immunofluorescence and Quantitative Microscopy

To visualize the expression of mScarlet and MBP-tagged SARS-CoV-2 Nsp1 (WT, CT, NT, Nsp1-linker1 and Nsp1-linker2), transfected HeLa cells were fixed with 4% formaldehyde for 10 minutes at room temperature, permeabilized with PBS + 0.25% Triton X-100, and blocked with 3% BSA. Cells were stained with mouse anti-MBP (NEB, 1:500), followed by washing and subsequent incubation with goat anti-mouse IgG, Alexa Fluor 488 (1:1000, Invitrogen) as well as Hoechst 33342 counterstain (Immunochemistry). Samples were imaged using a Nikon Eclipse Ts2R microscope or an ImageXpress Micro Confocal (Molecular Devices) with a 20X objective (S Plan Fluor, NA=0.45) using DAPI, GFP, and TRITC filter sets. Image data were analyzed and quantified using Image J (NIH) or MetaXpress software (Molecular Devices).

### Statistics

All of the experiments were independently performed in triplicate. The data were presented as mean ± SEM, except where noted otherwise. All graphs were plotted and analyzed with GraphPad Prism 5 software. p > 0.05 was considered statistically not significant, and the following denotations were used: **p < 0.001 and *p < 0.05.

### Data Availability

The cryo-EM maps included in this study have been deposited in the Electron Microscopy Data Bank with the accession code `EMD-22681`. The atomic coordinates have been deposited in the Protein Data Bank with the accession code `7K5I`.

## ACKNOWLEDGMENTS

The cryo-EM data collection was carried out by Thomas J. Edwards at the National Cancer Institute’s National Cryo-EM Facility, Frederick National Laboratory for Cancer Research under contract HSSN261200800001E. L.W. was supported by funding from an NIH T32 grant (5T32AI007512-34). T.-M.F. was supported by funding from an NIH T32 grant (5T32HL066987-18 to L.E.S.) and by start-up funds from the Ohio State University Comprehensive Cancer Center.

---

**Figure S1.**
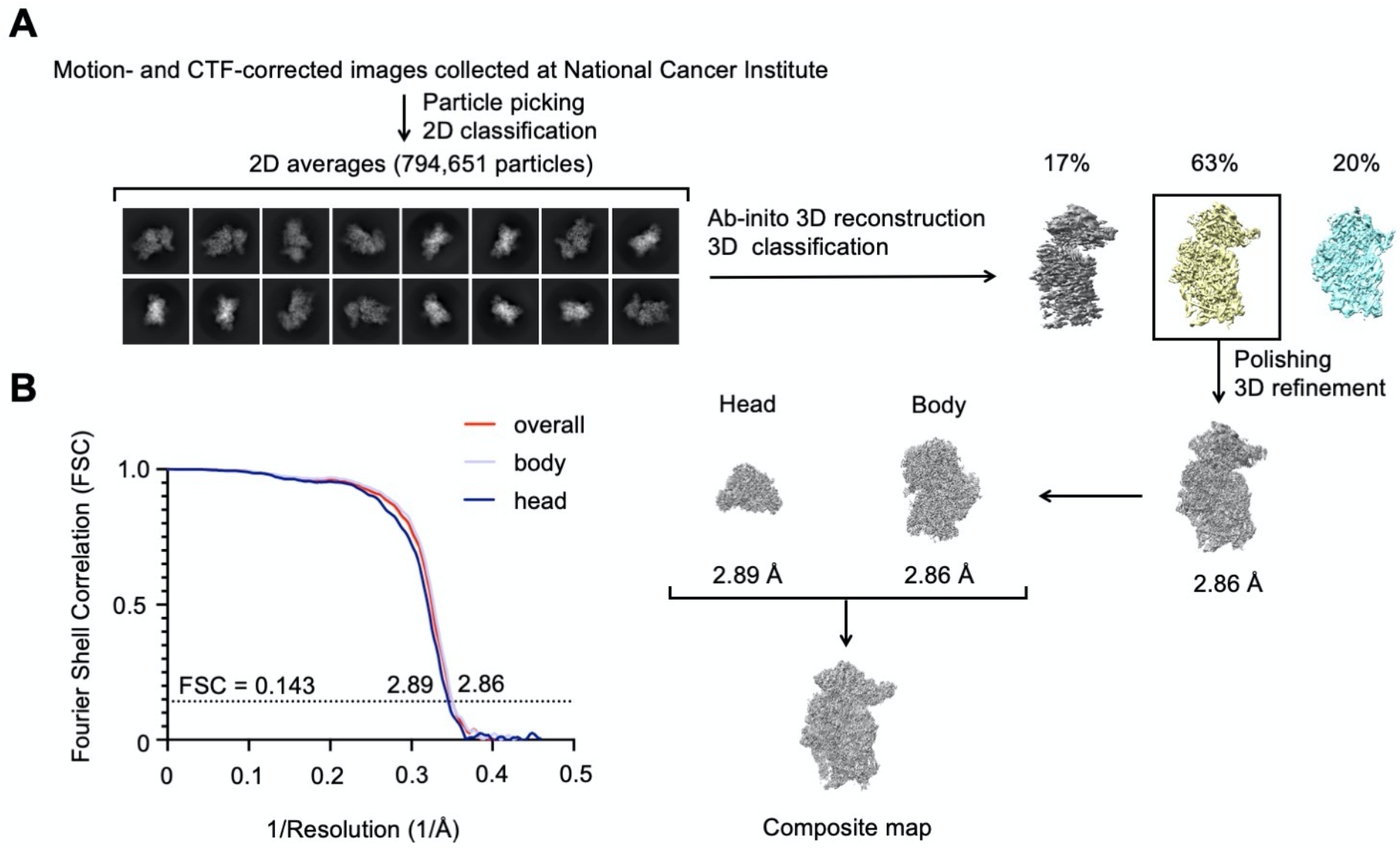
Reconstruction of SARS-CoV-2 Nsp1 bound to human 40S ribosomal subunit. (A) Workflow of 3D reconstruction. The selected map is boxed in rectangles. (B) Fourier shell correlation (FSC) curves of 3D reconstructed complex of SARS-CoV-2 Nsp1 and human 40S ribosomal subunit.

